# Optical photothermal infrared imaging of fatty acid esterification in the ER of living cells

**DOI:** 10.64898/2025.12.09.693044

**Authors:** Hannah B. Castillo, Caitlin M. Davis

**Affiliations:** Department of Chemistry, Yale University, New Haven, Connecticut, 06511, USA

## Abstract

Lipotoxicity is an accumulation of lipids that leads to cell death and metabolic disease. Saturated fatty acids are more likely to cause lipotoxicity, however, the mechanism remains unclear due to challenges visualizing reactions in live cells. Here, we use optical photothermal infrared (OPTIR) microspectroscopy to investigate palmitic acid (PA) metabolism in hepatocytes with sub-micron spatial resolution. Upon PA feeding, we discover a time-dependent ester carbonyl stretch localized to the ER and lipid droplet-ER contact sites of lipid droplets with abnormal morphology. This stretch is assigned to diacylglycerol intermediates in the glycerol-3-phosphate pathway. C-D stretches of deuterated PA provide complementary molecular details, supporting a model whereby PA acyl chain packing in the ER reduces enzyme diffusion slowing PA metabolism. Our results provide a deeper understanding of how phase changes induced by high melting temperature fatty acids and their metabolites change ER chemistry as well as provides a tool for detecting chemical and environmental changes associated with lipotoxicity in live cells.

**Teaser:** Sub-micron IR imaging of palmitic acid metabolism in live cells reveals diacylglycerol buildup and gel phase changes in the ER.

## Introduction

Metabolic diseases such as obesity, diabetes, and metabolic dysfunction-associated steatotic liver disease (MASLD), formerly known as non-alcoholic fatty liver disease, are a major health crisis (*1*). These conditions are widespread; with a worldwide prevalence over 30%, MASLD is the most common chronic liver disease (*2*). In these diseases, lipid-induced toxicity, also known as lipotoxicity, plays a major role in their onset and development (*1*). Excess lipid accumulation in the liver, termed steatosis, can lead to even more severe conditions, such as cirrhosis and liver cancer (*1*). A key cause of steatosis is fatty acid overload, when free fatty acid (FFA) influx and synthesis exceeds the liver’s ability to use FFAs as energy or store them in the form of triacylglycerols (TAGs) (*1*, *3*). While FFAs and their metabolites are essential for intracellular signaling and energy homeostasis, an overload of lipids can result in oxidative stress and cell death (*3*). Although these cell stress responses are well documented, due to difficulties directly visualizing metabolic species, the sub-cellular organization and molecular level details of the underlying chemical conversions that contribute to lipotoxicity are not well understood.

Lipid droplets (LDs) protect against lipotoxicity, serving as depots for neutral lipids. Ubiquitous in cells, LDs are composed of a core of neutral lipids bound by a phospholipid monolayer (*4*, *5*). Once regarded simply as storage pools of lipids, recent work has shown that LDs are dynamic organelle that regulate the flux of lipids through periods of growth and consumption (*4*). This process closely reflects the metabolic state of the cell, as the LD will sequester potentially toxic lipids in periods of fatty acid overload and release them when the cell is in need of energy (*5*). LDs are closely associated with the endoplasmic reticulum (ER), where the majority of hepatic TAGs are synthesized via the glycerol-3-phosphate (G3P) pathway (**Fig. 1**) (*4*, *6*). Precursors to this pathway include both exogenous fatty acids (FAs) taken up by the cell through diffusion or fatty acid transporter proteins (FATP) as well as *de novo* lipogenesis (DNL) derived FAs. Once in the cell, FAs must be activated by conjugation to coenzyme A (CoA) via long-chain acyl-CoA synthetases (ACSL), forming fatty acyl-CoA molecules (*6*). These molecules then undergo stepwise esterification to G3P by multiple enzymes in the ER membrane, ultimately forming TAGs (*7–9*). TAGs are deposited within the leaflets of the ER bilayer until they reach a critical concentration, resulting in phase separation and coalescing into a lipid lens that expands and eventually buds from the ER membrane in the form of a lipid droplet (*4*, *5*, *10*).

**Fig. 1.**
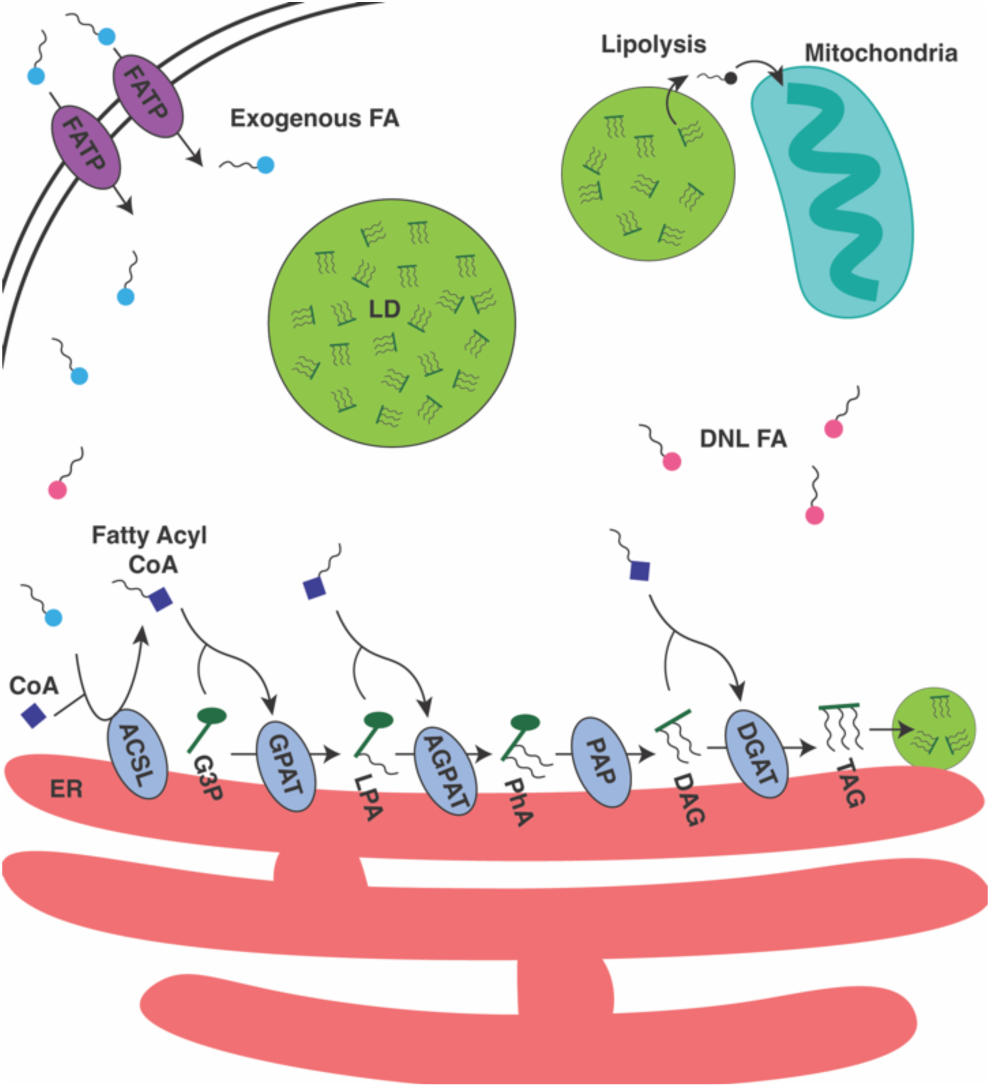
Schematic of triacylglycerol synthesis via the G3P pathway. FATP, fatty acid transporter protein; ACSL, Acyl-CoA synthetase; G3P, glycerol-3-phosphate; GPAT, G3P acyltransferase; LPA, lysophosphatidic acid; AGPAT, acylglycerol-3-phosphate acyltransferase; PhA, phosphatidic acid; PAP, phosphatidate phosphohydrolase; DAG, diacylglycerol; DGAT, diacylglycerol transferase; TAG, triacylglycerol; LD, lipid droplet; ER, endoplasmic reticulum.

While LDs serve as an excellent defense against lipotoxicity, increased levels of exogenous FAs can still lead to steatosis and have been shown to alter this carefully regulated pathway (*11*). Furthermore, saturated and unsaturated FAs have differential effects on cells. Excess levels of palmitic acid (PA), a saturated FA, induces inflammation, ER stress, and ultimately cell death (*12–14*). Conversely, excess oleic acid (OA), an unsaturated FA, while inducing steatosis, does not cause the same toxic effects and even suppresses PA-induced apoptosis in cells (*13*, *14*). These effects are also evidenced in the resulting LD morphology. While both FAs cause an increase in LD size in comparison to control conditions, LDs resulting from OA feeding are often significantly larger than LDs in PA-fed cells. Meanwhile, PA-feeding causes a larger increase in the number of LDs per cell than OA-feeding (*14*, *15*). This suggests that PA and OA trigger a spatial regulation of DNL involving LD morphology. While it is known that various FAs conversely contribute to cell fates and liver disease, missing is a mechanistic understanding of how such disruptors affect localized metabolism at the sub-cellular level. Several theories have emerged. One theory is that PA’s toxicity is due to its lower incorporation into LDs, changing the ratio of esterified to toxic unesterified FAs (*13*, *16*). Others attribute PA’s toxicity to downregulation of TAG synthesis enzymes contributing to metabolic dysfunction (*17*). Thus, investigating PA localization and metabolism at a subcellular level is essential for understanding its role in disease. However, traditional methods of tracking PA headgroup chemistry lack spatial resolution or use fixatives that can perturb localization. Further, while Raman imaging techniques can track the acyl tail they lack selectivity for chemistry at the FA headgroup.

Here, we leverage the sub-micron resolution of optical photothermal infrared (OPTIR) imaging to track PA metabolism in live hepatocytes (**Fig. 2A**). We have previously used OPTIR to track DNL and to monitor the effect of OA on DNL (*11*, *18*). In this work, we feed deuterated palmitic acid (PA-d_31_, ^2^H PA) to the hepatocyte-derived cell line Huh-7 and use C-D stretches and ester carbonyl shifts to monitor FA uptake, metabolism, and incorporation into LDs. Our results support both the theory that PA alters storage of FAs in LDs and downregulates TAG synthesis. Compared to OA, we observe slowed PA metabolism in the ER and at ER-LD contacts near LDs with abnormal morphology. We find that the two effects are coupled to phase changes in the ER that induce ER stress. Using the C-D stretches of ^2^H PA we demonstrate that upon PA feeding, TAG precursors in the G3P pathway enter a rigid gel phase by tightly packing their acyl chains. Using the C=O stretch of ^2^H PA we assign the TAG precursor to DAG. We conclude that acyl chain packing slows metabolism, leading to the buildup of DAGs in the ER, which in turn inhibits the ability of LDs to properly emerge from the ER, giving them an abnormal morphology. Azido PA, which packs less tightly than PA due to the azide, did not exhibit any of these effects. Similarly, unsaturated FAs like OA that have lower melting temperatures have been found to rescue cells from PA-induced apoptosis. Overall, this work sheds new light on the toxic origins of PA and other saturated fatty acids, as well as establishes new tools for understanding PA chemistry in the ER and the function of LDs in lipotoxicity.

**Fig. 2.**
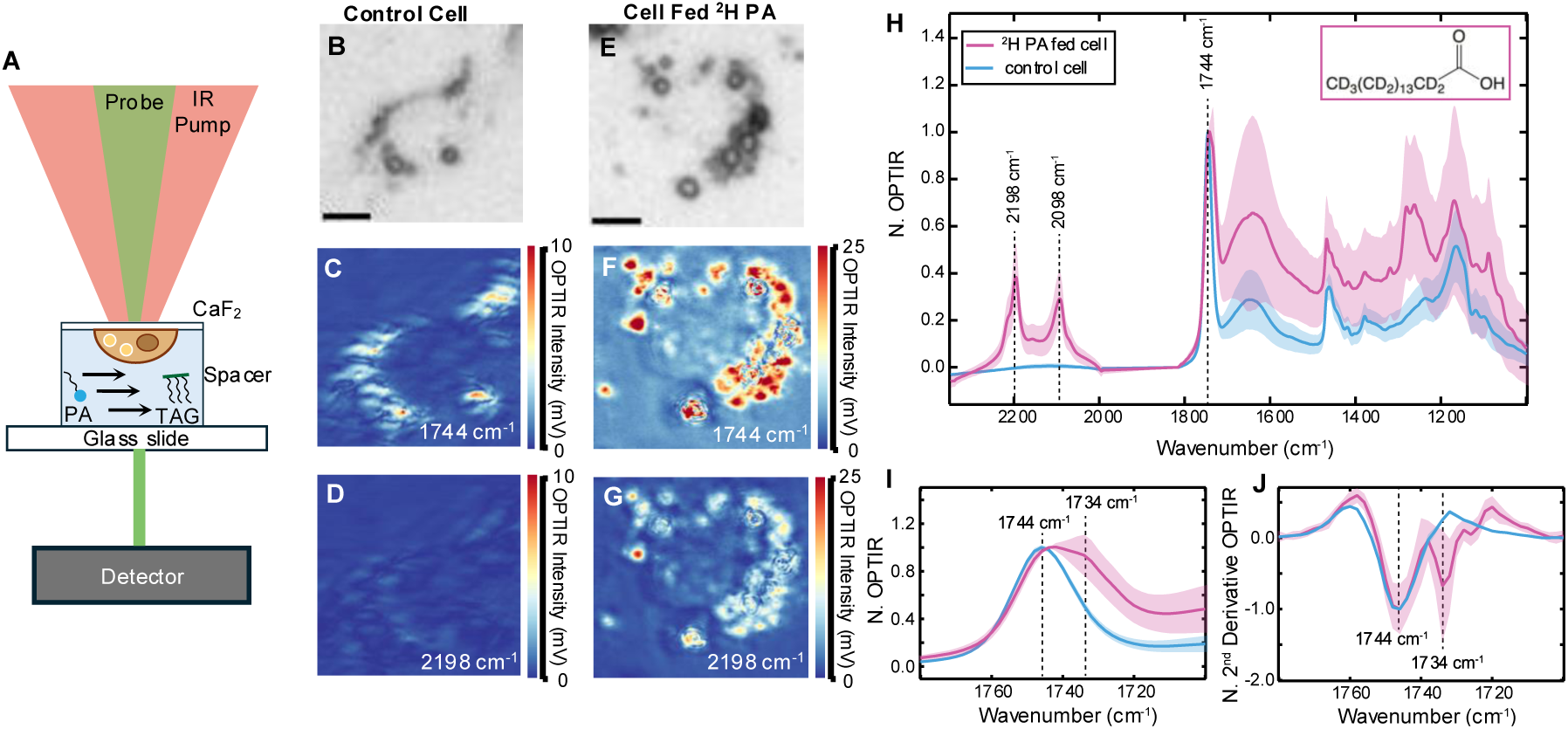
Huh-7 cells fed ^2^H PA exhibit an unanticipated shoulder off the C=O ester carbonyl stretch of TAGs/CEs. (**A**) Schematic of live cell imaging using OPTIR. (**B**) Brightfield of a live control Huh-7 cell (**C**) Single wavenumber image of the control cell collected at the C=O lipid carbonyl band (1744 cm^-1^) (**D**) Single wavenumber image of the control cell collected at the *v_as_* C-D_2_ stretching mode (2198 cm^-1^) (**E**) Brightfield of a live Huh-7 cell 24 hours after feeding 60 µM of ^2^H PA conjugated to BSA at a 2:1 ratio (**F**) Single wavenumber image of the ^2^H PA fed cell collected at the C=O lipid carbonyl band (1744 cm^-1^) (**G**) Single wavenumber image of the ^2^H PA fed cell collected at the *v_as_* C-D_2_ stretching mode (2198 cm^-1^) (**H**) Average normalized OPTIR spectrum collected in LDs of one control Huh-7 cell (blue) and six ^2^H PA-fed Huh-7 cells. PA-fed cells were imaged 24 hours after feeding 60 µM of ^2^H PA conjugated to BSA at a 2:1 ratio (pink). Shading is one standard deviation of the average normalized spectrum. The dominant C=O peak at 1744 cm^-1^, attributed to the ester carbonyl of TAGs/CEs, confirms that both cells are imaged on LDs. Only the cell fed ^2^H PA exhibits C-D_2_ asymmetric and symmetric stretches at 2198 cm^-1^ and 2098 cm^-1^, respectively. (**I**) A closeup of the C=O spectral region from 1700 to 1780 cm^-1^. An unanticipated shoulder at 1734 cm^-1^ is observed only in ^2^H PA fed cells. (**J**) A second derivative of the C=O region further resolves the shoulder at 1734 cm^-1^.

## Results

### Palmitic acid feeding reveals an unanticipated spectral feature at 1734 cm^-1^

To observe how PA is taken up and stored by the cell, we fed cells 60 µM ^2^H PA conjugated to BSA at a 2:1 ratio, in line with physiological free fatty acid concentrations that fluctuate from submillimolar to millimolar (*19*, *20*). Deuteration of the C-H bonds in PA shifts the frequency of the asymmetric and symmetric C-H_2_ stretching modes from 2916 cm^-1^ and 2850 cm^-1^ to 2198 cm^-1^ and 2098 cm^-1^, respectively (**Fig. S1**). These C-D_2_ stretches lie in the “cell silent region”, enabling detection of exogenously obtained PA without interfering signals from endogenously synthesized PA or other cellular molecules. Importantly, the ^2^H does not alter the structure of PA and, therefore, it accurately mimics PA. Further, the carbon chains are preserved as PA is metabolized, primarily through esterification and storage in LDs, allowing assignment of any species with these peaks as a derivative of ^2^H PA. Free PA is generally not observable outside of LDs due to the detection limit of the instrument (**Fig. S2**), in line with reported detection limits of similar instruments (*11*, *21*).

As most FAs end up esterified in LDs, initial spectra were collected on LDs of live Huh-7 cells 24 hours after feeding. Single wavenumber images were collected at frequencies attributed to ^2^H lipids to confirm the uptake of ^2^H PA. Control cells only fed BSA (vehicle control, **Fig. 2B**) only show signal in IR images collected at 1744 cm^-1^, originating from the ester carbonyl stretch of TAGs and cholesteryl esters (CEs) in the LDs (*22*) (**Fig. 2C**). No signal is seen in IR images collected at 2198 cm^-1^, the asymmetric C-D_2_ stretch that would indicate a ^2^H lipid (**Fig. 2D**). Cells fed ^2^H PA (**Fig. 2E**) show signal in IR images collected at both 1744 cm^-1^ and 2198 cm^-1^ that overlap well with each other as well as the LDs in the brightfield, indicating that the ^2^H PA has been taken up by the cell, esterified, and stored in LDs (**Fig. 2F-G**). This is also confirmed by the strong C-D stretches in single spectra collected on LDs of ^2^H PA-fed cells (**Fig. 2H**). Intriguingly, a 1734 cm^-1^ shoulder off this ester carbonyl peak (**Fig. 2I**) was observed in numerous PA-fed spectra. A second derivative further resolves this peak from the nearby 1744 cm^-1^ band (**Fig. 2J**). This spectral feature is only found in lipid-rich regions of cells fed ^2^H PA and is always associated with C-D stretches, suggesting that it is a derivative of PA and therefore a lipid. To confirm that the shoulder is not an optical artifact derived from the deuterium labels on the PA, unlabeled PA was fed to cells at the same concentration and the same shoulder was observed (**Fig. S3**).

### Timelapse of the 1734 cm-1 peak suggests a metabolic intermediate

We hypothesize that the 1734 cm^-1^ shoulder arises from a metabolic intermediate of PA in the G3P pathway that generates TAGs. If true, there would be a temporal component to the C=O stretch as the ^2^H PA is processed and not refed. To test this hypothesis, Huh-7 cells were incubated with ^2^H PA and monitored by OPTIR approximately every three hours for 72 hours. An average of 22 spectra were collected in LDs from approximately five cells per time point (total n=276) and each cell’s corresponding spectra averaged. This revealed a temporal trend to the 1734 cm^-1^ shoulder.

Throughout 72 hours, the LDs of cells grew with increasing time (**Fig. 3A-F**). For the first 12 hours, the 1744 cm^-1^ ester carbonyl peak of TAGs/CEs dominates the lipid region in the spectra. The peak at 1734 cm^-1^ first emerges as a small shoulder off the 1744 cm^-1^ peak around 12 hours after initial feeding (**Fig. 3G**). There was cell to cell heterogeneity in the exact time post-feeding that the shoulder presented, consistent with our past work that lipid metabolism is highly heterogeneous between cells (*11*, *18*). Over the next few hours, the intensity of the 1734 cm^-1^ band slowly increased until it was the dominating species in the ester carbonyl region. Between 27 and 54 hours the 1734 cm^-1^ peak was visible in almost all investigated cells. After which, it slowly merges back into the 1744 cm^-1^ peak between 52 and 70 hours after feeding. By 69 hours after feeding, the 1734 cm^-1^ peak is undetectable **Fig. 3G**). Taking the second derivative of the averaged spectra confirms this trend (**Fig. 3H**), with the 1734 cm^-1^ peak appearing around 12 hours post-feeding, growing and predominating over the 1744 cm^-1^ stretch, and shrinking until it is indistinguishable from the 1744 cm^-1^ triacylglycerol peak by 69 hours post-feeding.

**Fig. 3.**
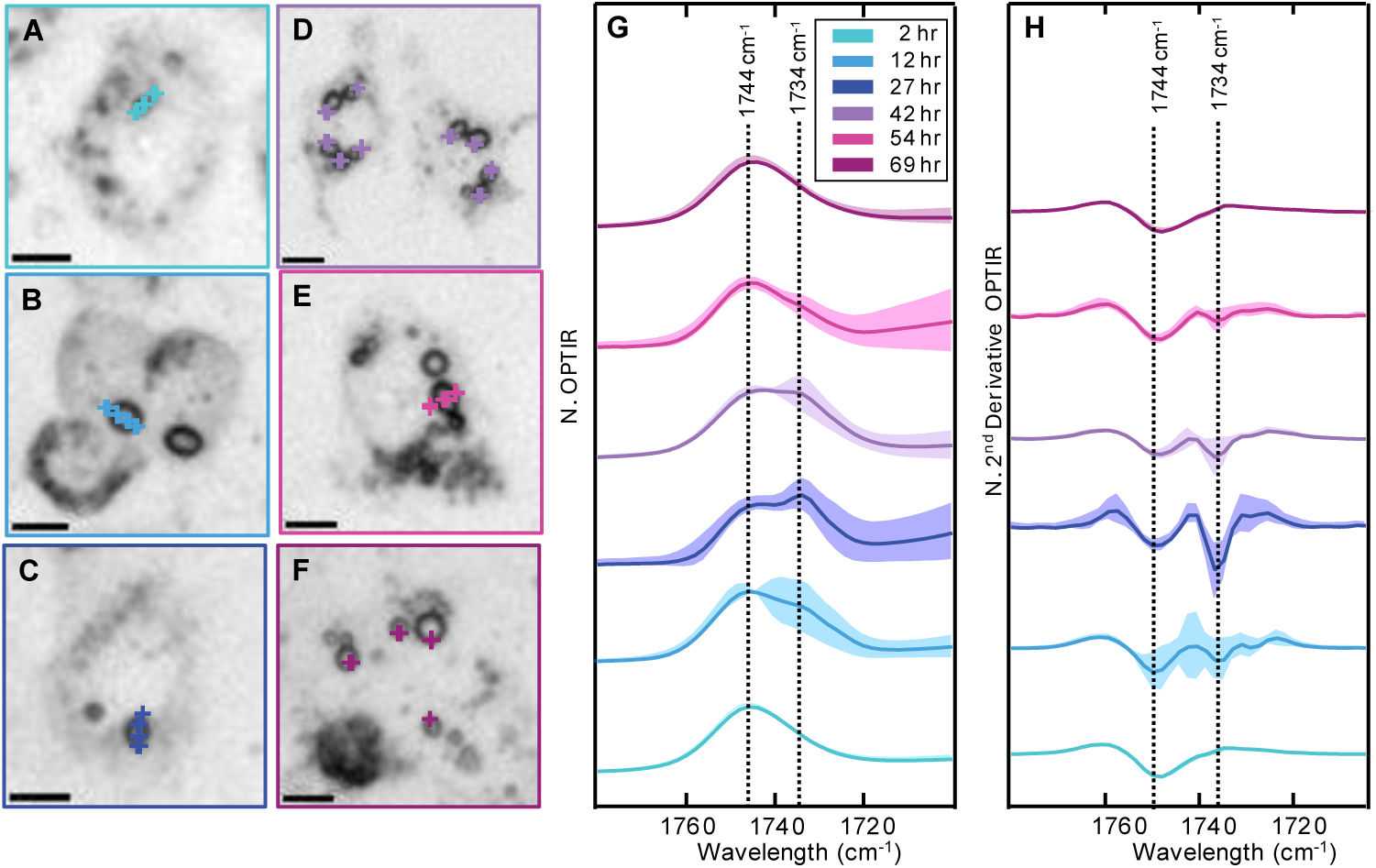
Timelapse of the ester carbonyl stretch region from 2 to 70 hours post feeding 60 µM ^2^H PA conjugated to BSA at a 2:1 ratio. (**A-F**) Brightfield images of Huh-7 cells fed at 2, 12, 27, 42, 54, and 69 hours, respectively. Colored crosses indicate where corresponding spectra was collected and is plotted in (G). (**G**) Average normalized IR spectra with standard deviation of Huh-7 cells fed ^2^H PA at 2, 12, 27, 42, 54, and 69 hours. Each time point represents the average spectra from the points labeled in (A-F). Shading is one standard deviation of the average normalized spectrum. (**H**) Second derivative of the average spectra in (G) with corresponding standard deviations.

This data is consistent with our hypothesis, that the unknown lipid peak is a metabolic intermediate that is built up to a concentration detectable via OPTIR over the first 27 hours, and then processed to a concentration undetectable via OPTIR over the following 42 hours. Further, from 12 to 27 hours, the second derivative reveals an additional peak at 1726 cm^-1^ that may indicate a low population of another metabolic intermediate (**Fig. 3H**).

### Intermediate buildup appears in the ER and at ER-LD contacts

Brightfield images reveal changes in the morphology of ^2^H PA-fed cells. LDs in cells fed ^2^H PA increased in both size and number compared to control cells (**Fig. 4A**). This observation agrees with other studies that report PA induces an increase in LD size and number (*14*, *23*). In addition to this, LD morphology is altered; many LDs in PA fed cells are oblong shape (**Fig. S4A-B**). These oblong LDs tend to have strong 1734 cm^-1^ signal and are only observed when feeding PA, not in control conditions nor when feeding unsaturated FAs such as OA (**Fig. S4C-F)**. Abnormal LD morphology in combination with the hypothesis that the 1734 cm^-1^ intermediate species is from TAG synthesis suggests that the buildup of these intermediates affects how LDs bud from the ER.

**Fig. 4.**
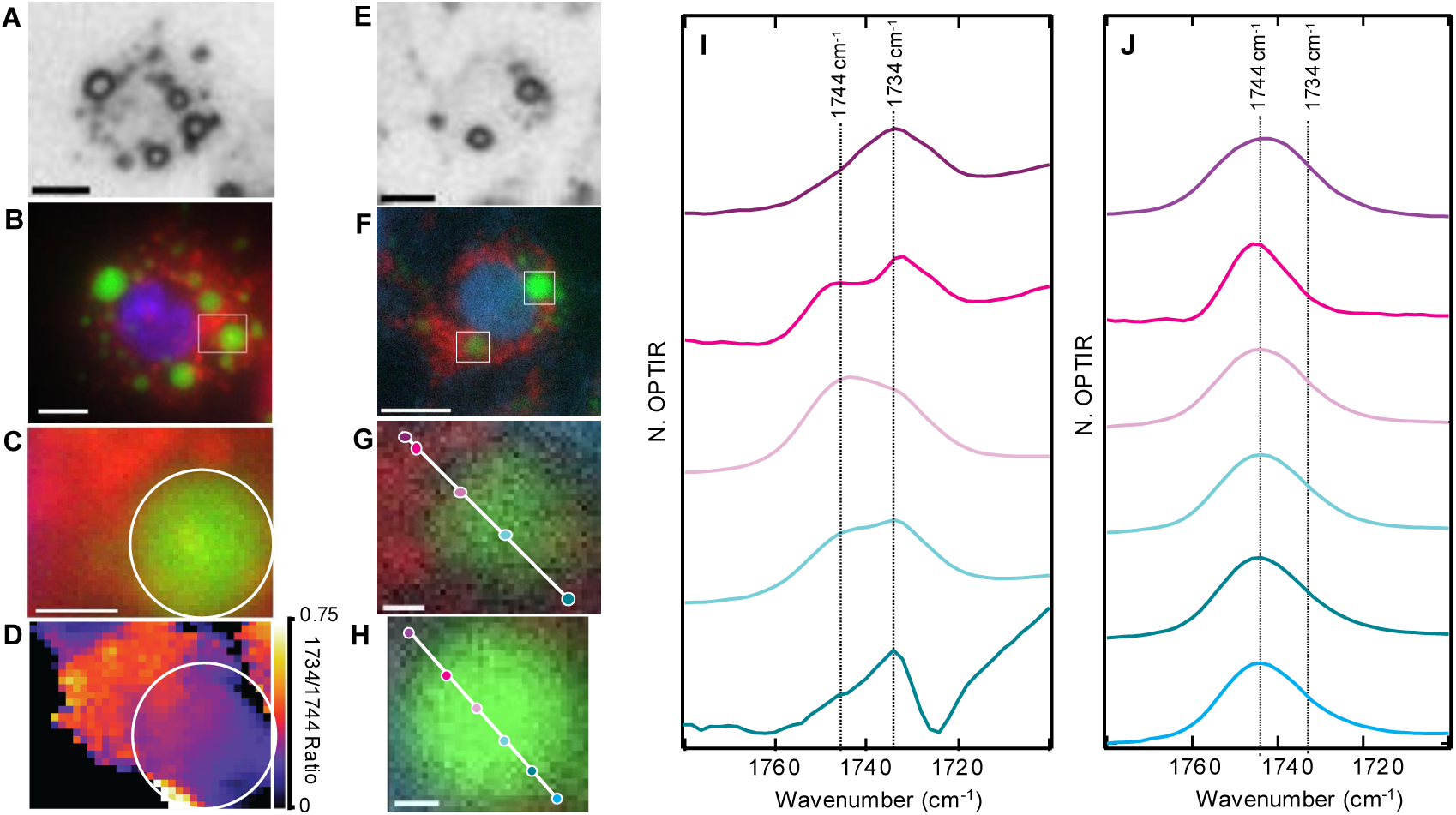
Fluorescence and IR images demonstrate that PA intermediates build up in the ER and at ER-lipid contact sites. (**A**) Brightfield image of a live Huh-7 cell 24 hours after feeding with 120 µM of ^2^H PA conjugated to BSA at a 2:1 ratio. Scale bar is 10 µm. (**B**) Fluorescence image of the cell in (A) stained for the ER (red), nucleus (blue), and LD (green). A white box indicates where a hyperspectral image was collected. Scale bar is 10 µm. (**C**) Fluorescence signal in the region where the hyperspectral image was collected. The lipid droplet is outlined in white. Scale bar is 1 µm. (**D**) Ratio of the 1734 cm^-1^ band to the 1744 cm^-1^ band in the hyperspectral image with the lipid droplet outlined in white. **(E)** Brightfield of a Huh-7 cell 24 hours after feeding with 120 µM of ^2^H PA conjugated to BSA at a 2:1 ratio. (**F**) Fluorescence image of the cell in (E) stained for the ER (red), LD (green), and nucleus (blue). Two white boxes indicate the LDs that were studied. Scale bar is 10 µm. **(G)** Expansion of a LD with high ER fluorescence from (F). A line scan was collected across this LD along indicated points. Scale bar is 1 µm. **(H)** Expansion of a LD with low ER fluorescence from (F). A line scan was collected across this LD along indicated points. Scale bar is 1 µm. **(I)** Spectra corresponding to the points in (G) where a linescan was collected across the LD. Ester carbonyl bands at 1744 cm^-1^ and 1734 cm^-1^ are marked. **(J)** Spectra corresponding to the points in (H) where a linescan was collected across the LD. Ester carbonyl bands 1744 cm^-1^ and 1734 cm^-1^ are marked. All images collected on the OPTIR.

LDs have many contact points with other organelle, including the ER, mitochondria, and peroxisomes (*4*, *24*). These contact points form metabolic hubs imperative to FA homeostasis and facilitate FA trafficking, contributing to LD growth and degradation (*4*, *24*). We focus on the ER because TAG synthesis and LD nucleation occur in the ER. As we believe the 1734 cm^-1^ peak is a ^2^H PA metabolite, it should increase in intensity when the concentration of ^2^H PA fed to cells is increased. Thus, to better resolve the shoulder, Huh-7 cells were incubated with ^2^H PA at a slightly higher physiological concentration of 120 µM conjugated to BSA at a 2:1 ratio. After 24 hours, cells were fluorescently stained for the nucleus, ER, and LD with Hoechst nuclear stain (blue), ER-Red (red), and LipiSpot (green), respectively. Cells were then imaged live on the OPTIR. Cells with multiple, large LDs were chosen for analysis (**Fig. 4A**).

To determine if the 1734 cm^-1^ peak is associated with the ER and/or spatially confined to the LDs, hyperspectral images were collected across LDs in regions with high ER fluorescence where it is likely that the LD has contact sites with the ER (**Fig. 4B-C**). Hyperspectral maps are three-dimensional images comprised of OPTIR spectra from 1600 to 2348 cm^-1^ collected at every pixel within the area of interest. To reduce noise from lipid free areas that may be amplified when creating ratio images, only regions with sufficient 1744 cm^-1^ signal were analyzed (see Materials and Methods). From a hyperspectral map, a ratio image of the 1734 cm^-1^ peak to the 1744 cm^-1^ peak reveals the spatial distribution of the metabolic intermediate relative to TAGs/CEs. Higher 1734 cm^-1^/1744 cm^-1^ ratios are found both at the edge of LDs bordering the ER and in the ER itself (**Fig. 4D**). This indicates that the metabolic intermediate is concentrated in the ER and at LD-ER contact sites.

To further confirm that the intermediate is arising from chemistry in the ER, a cell with both a LD in an ER rich region and a LD in an ER free region was investigated (**Fig. 4E-F**). Linescans were collected over both LDs (**Fig. 4G-H**). Like the droplet in **Fig. 4A**, the LD with potential to make ER contacts (**Fig. 4G**) had regions of increased 1734 cm^-1^ intensity at the LD edge bordering the ER At the edges of the LD the 1734 cm^-1^ band dominates, while the 1744 cm^-1^ band dominates in the center of the LD (**Fig. 4I**). In contrast, the LD in an area without potential to make significant ER contacts (**Fig. 4H**) did not have any regions with significant 1734 cm^-1^ signal (**Fig. 4J**), indicating that there are no metabolic intermediates building up in or around that LD. This further highlights the heterogeneity of the LDs within cells, as some exhibit this 1734 cm^-1^ lipid species and some do not. This trend was confirmed through complementary fluorescence and infrared analysis of 8 cells; only LDs in ER rich regions have the 1734 cm^-1^ signal, and thus metabolic intermediate buildup. As TAG synthesis occurs in the ER and LDs are known to have multiple contacts with the ER even after budding, we hypothesize that the 1734 cm^-1^ lipid species is a precursor of TAG synthesis arising from PA feeding.

### DAGs accumulate in Huh-7 cells upon PA feeding

Previous work has shown that lipid metabolism is affected by PA feeding (*17*, *25*, *26*). Cells fed PA tend to have large increases of glycerolipids in comparison to control conditions. Specifically, lipodomics find that large amounts of disaturated DAGs accumulate (*25*, *26*). We hypothesize that the 1734 cm^-1^ lipid species is a disaturated DAG building up to a concentration that is observable by OPTIR.

TAG synthesis by the G3P pathway includes five primary lipid species: a free fatty acid, LPA, PhA, DAG and TAG (**Fig. 5A**). To assign the 1734 cm^-1^ peak, we compare it to the OPTIR spectra of these five lipid species. As we established the unassigned dominant intermediate arises from PA feeding, the acyl chains of all TAG precursors analyzed were solely comprised of PA. IR spectra for palmitic acid, palmitoyl lysophosphatidic acid, 1,2-dipalmitoyl-sn-glycero-3-phosphate, 1,2-dipalmitoyl-sn-glycerol, and tripalmitin were collected on the OPTIR. However, the vibrational frequency of a bond, especially carbonyls, is dependent on the local electric field as well as nearby hydrogen bonds (*27*). Thus, the spectrum of each lipid species may differ slightly depending on the solvent. Previous work has shown that the ER has a polarity between chloroform and dichloromethane (DCM) (*28*). Therefore, to replicate the polarity of the ER, all five precursors were dissolved in chloroform at a concentration of 25 mM (**Fig. 5B**). The solubility of PhA and LPA in DCM was below the detection limit of the OPTIR, therefore, only PA, DAG, and TAG were compared in DCM (**Fig. S5**). Comparing the OPTIR spectra in chloroform, the carboxylic acid carbonyl of PA is at 1709 cm^-1^, while the rest of the precursors have ester carbonyls with peaks that are in the range of 1730-1740 cm^-1^. As anticipated, the bandwidth and peak position of the ester carbonyl of DAGs in chloroform best matches the *in cellulo* peak at 1734 cm^-1^. In DCM, DAG appears at 1736 cm^-1^, also closely matching the 1734 cm^-1^ peak. Thus, we assign the 1734 cm^-1^ peak to DAGs.

**Fig. 5.**
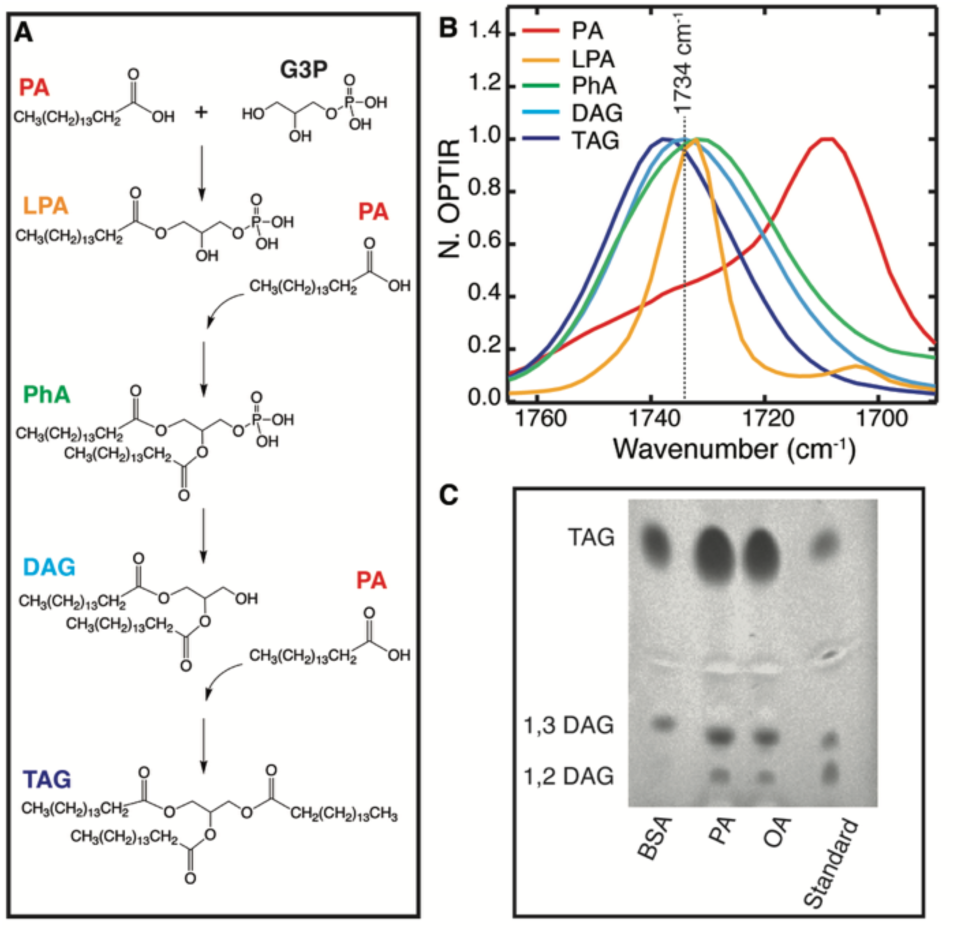
Assignment of 1734 cm^-1^ to DAGs. (**A**) TAG synthesis via the G3P pathway with PA as the FA chains. (**B**) OPTIR spectra of TAG intermediates dissolved in chloroform at 25 mM. (**C**) TLC plate of lipids isolated from PA-fed, OA-fed, and control cells. Cells were fed 60 µM of BSA-conjugated PA or OA or BSA alone (control) and incubated for 24 hours. Tripalmitin (TAG), 1,3 dipalmitin (1,3 DAG), and 1,2 dipalmitin (1,2 DAG) were used as standards.

To further confirm the DAG assignment the cell’s lipid profile was assessed using thin layer chromatography (TLC). Cells were incubated with 60 µM of BSA-conjugated PA or OA or BSA alone (vehicle control) for 24 hours. Lipids were then isolated from the cell using the Bligh and Dyer lipid extraction technique and spotted onto TLC plates (**Fig. 5C**). In comparison to control cells fed BSA, PA- and OA-fed cells both produced large amounts of TAGs and 1,2 DAGs. This partially agrees with a previous study that used TLC to separate lipid extracts from cardiomyoblasts, however, in cardiomyoblasts PA led to the formation of significantly more DAGs than OA (*29*). In hepatocytes, the increase in DAGs upon PA-and OA-feeding is comparable (**Fig. 5C**). This is interesting as DAGs are only easily detected using OPTIR on PA-fed hepatocyte cells. We previously used the same OPTIR approach to monitor ^2^H OA feeding in hepatocytes and adipocytes, and no such TAG precursor peak had been observed (*11*). To further investigate, Huh-7 cells were fed 60 µM of ^2^H OA conjugated to BSA at a 2:1 ratio and then studied using OPTIR throughout 24-48 hours. Linescans were collected through large LDs. Surprisingly, a few LDs exhibited a shoulder at 1738 cm^-1^ off the 1744 cm^-1^ carbonyl stretch (**Fig. S6A**). Although this peak was found in OA-fed cells, it was far less common (**Fig. S6B**). Furthermore, no LDs with abnormal morphology were observed. This suggests that similar amounts of TAGs and DAGs are present in both PA- and OA-fed Huh-7 cells, but their spatial distributions and cell phenotypes differ. That DAGs are infrequently detected by OPTIR in OA-fed Huh-7 cells indicates that they are distributed across the cell such that the local concentration is below the detection limit of the OPTIR. Thus, another factor exists that is contributing to the significant local buildup of DAGs in the ER of PA-fed Huh-7 cells.

### ER rigidity is correlated with metabolic inhibition

A physical difference between OA and PA is their melting temperature. In general, saturated fats have higher melting temperatures than unsaturated fats because their straight acyl chains can be tightly packed. Indeed, the melting temperature of OA is ∼13 °C, while the melting temperature of PA is ∼63 °C (*30*). This trend is maintained through the five primary lipid species in TAG synthesis; the melting temperature of triolein is ∼5 °C while the melting temperature of tripalmitin ∼45-66 °C (*30*, *31*). Thus, at physiological temperatures OA and its TAG intermediates are a liquid whereas PA and its TAG intermediates are a solid. Mixtures of FAs have lower melting temperatures than pure FA (*32*), however, FA feeding artificially inflates the cellular distribution of the fed FA. We therefore hypothesize that PA and its TAG intermediates solidify in the ER, leading to ER stress and DAG accumulation.

To test this hypothesis, we investigate the C-D stretches of the acyl chains. As described above, differences in solvent polarity were considered in the assignment of the shifted lipid carbonyl peak. However, other physical properties of lipids can alter vibrational spectra, most notably the conformation and structure of the lipid (*33–35*). Lipids are polymorphic and under physiological conditions can undergo a lamellar gel to lamellar liquid-crystalline phase transition. The lamellar gel state is characterized by fully extended and rigid acyl chains in an all-*trans* conformation, resulting in ordered and tightly packed lipids (*33*). In contrast, the lamellar liquid-crystalline state is a disordered environment, with more hydrocarbon chains in the *gauche* conformation, allowing for increased dynamics of lipid molecules (*33*). Previous studies have demonstrated that the C-H vibrational modes can distinguish between these two states (*36*, *37*). The liquid crystalline phase is characterized by higher frequencies and broadening of the CH_2_ bands due to the increase in dynamics between acyl chains.

The C-D vibrational modes of Huh-7 cells were imaged by OPTIR 24 hours after ^2^H PA feeding. There is a stark difference between the symmetric and asymmetric CD_2_ stretches in spectra with and without the 1734 cm^-1^ lipid carbonyl peak. In an OPTIR spectrum taken directly on a LD, only one lipid carbonyl dominates at 1744 cm^-1^. Consistent with a liquid-crystalline phase, the corresponding symmetric and asymmetric CD_2_ stretches are located at 2098 cm^-1^ and 2200 cm^-1^, respectively (**Fig. 6A, bottom**). These are TAGs made with ^2^H PA acyl chains esterified and stored in LDs. When focused on the edge of a LD, the 1734 cm^-1^ peak dominates the lipid carbonyl region and the symmetric and asymmetric CD_2_ stretches redshift to 2090 cm^-1^ and 2194 cm^-1^, respectively (**Fig. 6A, top**). This is consistent with ^2^H PA acyl chains in a lamellar gel phase. Further, a small peak appears at 2158 cm^-1^, which can be assigned as Fermi resonance between the symmetric CD_2_ mode and the first overtone of the CD_2_ bending mode (**Fig. 6B**) (*36*, *38*, *39*). Fermi resonant interactions are sensitive to conformational changes of a molecule and have been used to monitor intermolecular interactions (*40–42*). This particular Fermi resonance has been reported to disappear during the lipid transition from the ordered lamellar gel phase to the liquid crystalline phase (*36*, *43*). Second derivative spectra confirm the appearance of the Fermi resonance as well as the shifting of the CD_2_ peaks (**Fig. 6C**). These results are reproducible across multiple spectra with and without the 1734 cm^-1^ peak. In most cases, the dominance of the 1734 cm^-1^ peak in the lipid carbonyl region is associated with a CD_2_ redshift and the appearance of the Fermi resonance. As previously established, these spectral features are most common at the edges of a LD that has strong contact points with the ER. Thus, as hypothesized, we conclude that it is lipids in the ER that are forming a lamellar gel state due to the incubation of PA which creates tighter packing of the acyl chains of TAG precursors in the ER. Most biomembranes, including the ER membrane, exist in the liquid crystalline state to allow for diffusion of biomolecules (*44*). We further speculate that enzymatic reactions necessary for TAG synthesis are diffusion limited in the lamellar gel state, slowing PA metabolism.

**Fig. 6.**
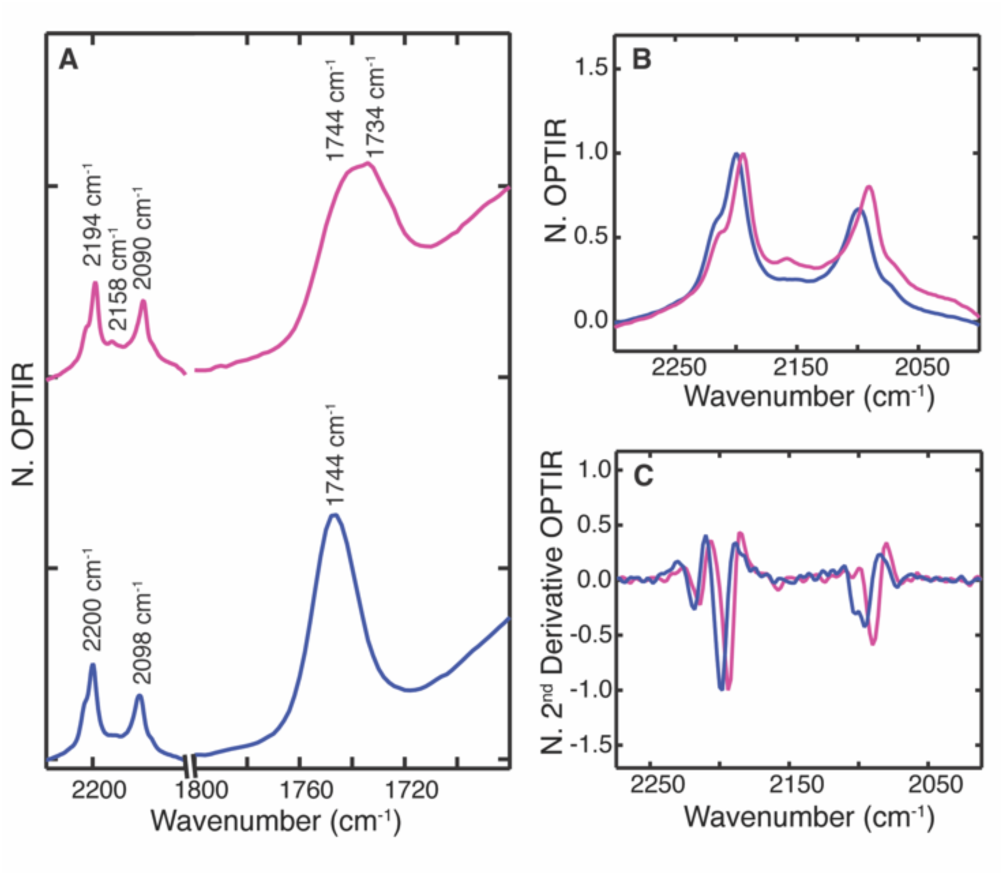
1734 cm^-1^ signal appears in an ordered lamellar gel environment. (**A**) A redshift of the asymmetric and symmetric CD_2_ stretches and the appearance of a Fermi resonance at 2158 cm^-1^ is correlated with the appearance of a 1734 cm^-1^ shoulder off the lipid carbonyl. Enlarged view of the C-D region and its corresponding second derivative spectra is shown in (**B**) and (**C**), respectively. Measurement of multiple cells and samples produced the same results. Spectra were normalized by prominent peak heights of the C-D region.

In addition to C-D stretches, azides are particularly sensitive to local environmental changes (*45*, *46*). To gain further insight into the environmental changes that occur in the ER upon PA build up, cells were fed 120 µM of azido PA. The azide peak appears at 2196 cm^-1^ in the cell silent region and is expected to redshift upon exposure to a nonpolar environment. However, no 1734 cm^-1^ lipid peak was found in any cells fed azido PA and the azide signal was only observed in LDs (**Fig. S7**). This suggests that azido PA TAG precursors do not build up and form a lamellar gel state in the ER like ^2^H PA and unlabeled PA. Unlike deuteration, which preserves the PA structure, incorporation of an azide slightly modifies the PA structure. Azido PA has a bulky azide group on the end of the hydrocarbon tail, which increases the area per molecule, reducing lipid thickness and restricting how closely molecules can pack together. Thus, our measurements suggest that azido PA has a melting point below PA and below physiological, 37 °C. Azide FAs have widely been used in vibrational studies, yet, to our knowledge, the 1734 cm^-1^ shoulder has never been reported (*47*, *48*). This presents a pitfall in the use of azido FAs as the azide modification affects FA metabolism. In addition, this observation confirms the correlation between the lamellar gel state and DAG accumulation.

To further investigate the extent of this effect, 120 µM of a 1:10 mixture of azido PA to unlabeled PA was fed to cells and imaged 24 hours after feeding. Similarly, no 1734 cm^-1^ lipid peak was found in any cells. This suggests that even low concentrations of azido PA disrupt lamellar gel state formation. It is well established that OA prevents PA-induced apoptosis. Ratios of OA to PA ratios as low as 1:25 have been recorded to have a protective effect against PA-induced apoptosis (*13*, *16*, *49*). Unsaturated long chain FA have lower melting temperatures and exist as liquids at room temperature. We hypothesize that the double bond of OA, similarly to the relatively bulky azide group of azido PA, disrupts formation of the lamellar gel state in the ER resulting in unrestricted movement in the ER, maintaining metabolic flux, and sequestering FAs into LDs to prevent apoptosis.

As reported above, a few Huh-7 cells fed ^2^H OA exhibited shifts in their ester carbonyl consistent with buildup of metabolic intermediates. To investigate whether this was correlated with a change in the state of the ER, 60 µM ^2^H OA was fed to Huh-7 cells that were monitored by OPTIR every three hours for 72 hours. As anticipated, the majority of ^2^H OA-fed cells do not exhibit any lipid carbonyl shoulders or shifts. However, as previously observed, a few cells do exhibit broadening and shifts of lipid carbonyl peaks and corresponding CD_2_ shifts, indicating that there is a buildup of metabolites and a phase change at the ER-LD contact sites (**Fig. S8**). Still, this shift was found less frequently in OA-fed cells compared to PA-fed cells. Between the timepoints of 1-55 hours, 12% of OA-fed cells exhibited a lipid carbonyl shift, compared to the 45% of PA-fed cells. Generally, hepatic LDs contain more TAGs than CEs (*50*, *51*). One possibility is that CEs can also undergo a phase transition, cholesteryl oleate and cholesteryl palmitate have melting temperatures above physiological, ∼44 °C and ∼76 °C, respectively (*52*, *53*). Thus, it stands that C-D shifts and DAG lipid carbonyl peaks are more common in PA-fed cells, as both palmitate TAG precursors and cholesteryl palmitate have higher than physiological melting temperatures. Only cholesteryl oleate has a higher than physiological melting temperature in OA-fed cells, resulting in less common C-D shifts. This suggests that not only are TAG precursors accumulating and forming a lamellar gel phase, but cholesteryl esters contribute to this as well.

## Discussion

Despite incredible advancements in understanding the roles LDs and PA play in metabolic disorders, many techniques cannot sufficiently answer the outstanding questions in the field. Lipidomics and biochemical assays have provided insight to how PA changes the FA profile but are lacking in spatial information essential to connect these effects to the cell at a sub-cellular level (*13*, *25*). Enzyme inhibition studies have spurred hypotheses on how PA disrupts metabolism, but look only at a particular metabolic step rather than the entire cellular system or trigger off-pathway compensations (*17*). Optical imaging is a promising approach to spatially resolve cellular metabolism. Techniques including magnetic resonance imaging and fluorescence imaging have made great progress in visualizing cellular metabolism and have contributed to the development of drugs and disease prognosis (*54*). However, these techniques have their own limitations, whether it be sensitivity, suitability for dynamic imaging, or use of bulky and perturbative probes.

Vibrational microspectroscopy coupled with vibrational probes has the potential to be an extremely powerful technique in the field of cell biology. Vibrational probes are small and biorthogonal, allowing for their incorporation and detection amongst other exogenous cellular molecules (*47*). Additionally, vibrational probes are environmentally sensitive providing information on the physical properties of the cell, which are difficult to obtain with conventional probes. As FAs are simple to label and easy for cells to uptake, many vibrational studies have monitored FA metabolism. Raman microspectroscopy has analyzed deuterated and alkyne-tagged PA’s effect on cells and whole organisms (*35*, *55*). Shen et al. used the Raman C-D vibrational mode along with fluorescent stains and HPLC-MS to monitor the formation of solid-like domain separation in the ER induced by ^2^H-PA feeding in HeLa cells (*56*). However, like other Raman studies, they focus on the acyl chains, as C-H bonds are particularly Raman active while carbonyl bonds are weak (*57*). This presents a disadvantage of Raman imaging, as it is not possible to directly observe the chemistry that occurs at the lipid headgroup during TAG synthesis.

IR is more sensitive to carbonyl bonds, and, with the recent development of sub-micron OPTIR, IR imaging now provides sufficient resolution to probe carbonyl metabolism at the sub-cellular level. Recent OPTIR studies used labeled PA but, like previous Raman studies, focus on the C-H and C-D regions. Park *et al.* monitored neutral lipid synthesis in live cells using ^2^H PA, but did not report any spectral shifts in the ester carbonyl region of TAGs/CEs (*21*). Shifts in the C-D region were attributed to heterogeneity of TAG acyl groups. While this could partially contribute to the broadening and shifts that we observe, the fermi resonance in our data is strong spectral evidence of a liquid crystalline to lamellar gel phase change. Teng *et al*. report FA desaturation of ^2^H PA through monitoring the vibration of the unsaturated C-D bond (*58*). They do not report C-D shifts, and we do not observe an unsaturated C-D bond vibration upon ^2^H PA feeding, but this may be due to a difference in cell lines. Bai *et al*. also monitored lipid synthesis over various model systems, but used azido-PA, which we discover does affect FA metabolism by disrupting the formation of a lamellar gel state (*48*). While these studies made significant contributions to the field, our work highlights the importance of using probes that do not alter FA metabolism and that can directly monitor where the chemistry occurs, at the headgroup of the lipid.

DAGs are involved in many cellular functions, mostly as signaling molecules and precursors to glycerolipids (*59*). As a signaling molecule, it is thought that DAGs block insulin signaling, resulting in insulin resistance, a symptom of type 2 diabetes and hepatic steatosis, both of which are often simultaneously observed in patients (*60*, *61*). Thus, buildup of DAGs has been implicated in hepatic diseases, but the extent of its impact on PA lipotoxicity is still debated. Our evidence suggests that DAGs are a necessary component to PA lipotoxicity, as it is the longest lived lipid intermediate in the G3P pathway (**Fig. 3**). Previous work has implicated DGAT1, which converts DAGs to TAGs, as a reason for DAG buildup and inhibition of TAG synthesis (*17*, *26*). Unsaturated FAs are thought to be superior substrates for TAG synthesis, while saturated FAs and DAGs are thought to be poor substrates that impair cellular flux, leading to accumulation of saturated lipid species (*5*, *26*). This aligns well with other findings that have found PA is less likely to be esterified into LD compared to OA, most likely due to TAG inhibition from saturated substrates (*12*, *13*, *26*). While our measurements agree that there are spatial differences in the esterification of OA and PA into LDs, we find that they result in a comparative buildup of TAG intermediates (**Fig. 5C**). We also find that both OA and PA have similar upstream effects on metabolic pathways. Consistent with our past OA studies, compared to control conditions, cells fed ^13^C glucose and ^2^H PA exhibit lower incorporation of ^13^C in TAGs/CEs (**Fig. S9**) which we attribute to downregulation of DNL-associated enzymes (*11*). Rather than saturated substrates inhibiting TAG enzymes in the G3P pathway, our measurements point towards an alternate, physical mechanism as the origin of DAG buildup in the ER.

In the ER, the C-D stretches of PA derived intermediates were consistent with a lamellar gel state. As saturated lipids can pack tighter together than unsaturated lipids, increased membrane order induces ER stress (*25*, *62*). Many studies have linked PA to increased ER stress and in turn, poor cellular health (*17*, *25*, *63*). This is due to altered lipid composition of the ER, which disrupts homeostasis and ultimately activates stress responses (*62*, *64*). Normally, to combat this, TAG synthesis enzymes relocalize from the ER to the budded LD surface via membrane tethers (*65*). However, upon elevated DAG levels, LD remain embedded in the ER. The conical shape of DAGs have a negative intrinsic curvature, which disfavors budding and promotes an embedded state of LDs (*4*, *66*). Our findings support this model. DAGs were only seen at the edges of LDs co-localized with the ER, suggesting that these LDs were still embedded in the ER. Further, the oval shaped LDs that were observed imply that there are issues with proper budding. These were also the LDs that were more likely to accumulate DAGs. Furthermore, as azido PA cannot be packed as tightly as PA, it did not induce DAG accumulation. Similar effects are seen upon OA feeding. Numerous studies have reported that as little as 1:25 OA can reverse negative effects of PA (*13*, *14*, *49*). OA likely disrupts the packing of PA, thereby reducing ER stress due to its lower melting temperature and increased enzyme diffusion. Thus, it is a physical change rather than direct enzyme inhibition that gives rise to the altered PA metabolism. Although there are still many questions about LD growth and function, our findings contribute a spatial understanding of the role PA chemistry and disaturated DAGs play in LD dynamics and morphology.

Using OPTIR, we have visualized DAG accumulation upon PA feeding through the emergence of a time-dependent ester carbonyl peak and monitored physical properties of the ER through C-D stretches in the acyl chains. To our knowledge, this is the first time TAG intermediates have been directly detected in live cells. Notably, this chemistry could be tracked entirely label free. In addition, OPTIR allowed us to monitor lamellar gel and liquid crystalline lipid states in the ER and LDs using shifts in the C-D region. Through these observations, we have established a mechanism by which PA induces lipotoxicity in the cell. As saturated PAs are imported to the ER for TAG synthesis, acyl chain packing of PA and associated intermediates induces a phase change to an ordered lamellar gel state that slows diffusion of enzymes necessary for TAG synthesis. ER stress occurs through the accumulation of disaturated DAGs and other lipid intermediates as well as LDs embedding in the ER. LD-organelle contact sites are disrupted as the LDs have not properly budded, interfering with lipid metabolism. This provides new insight to how PA induces lipotoxicity and establishes OPTIR as a highly effective technique to monitor lipid metabolism.

## Materials and Methods

### Materials

Unless otherwise specified, all materials were sourced from Sigma-Aldrich.

### FA conjugation

Fatty acid conjugation was performed as previously described using a protocol adapted by Shi *et al* (*11*, *47*). Briefly, in a 2 mL clear glass container (VWR, Radnor, PA), ^2^H palmitic acid (palmitic acid-d_31_, DLM 215, Cambridge Isotope Labs, Tewksbury, MA), or azido palmitic acid (CCT 1346, Vector Laboratories, Newark, CT) was dissolved in 29 mM NaOH and heated in a 70 °C water bath until clear. The fatty acid solution was then diluted to a total concentration of 3 mM with 150 mg/mL fatty-acid free BSA (MP Biomedicals, Santa Ana, CA) dissolved in DPBS (Thermo-Fischer, Waltham, MA). The solution was heated again at 37 °C in a water bath until clear and filtered with a 0.2 um sterile filter. Final ratios of palmitic acid to BSA were 2:1, in line with physiological ratios (*19*).

### Huh-7 cell culture

Huh-7 cells (a gift from the Yale Liver Center) were cultured as previously described (*11*). Briefly, cells were grown to 80% confluence in Dulbecco’s modified Eagle’s medium containing 4.5 g/L glucose and L-glutamine (DMEM, Corning, Corning, NY) supplemented with 10% fetal bovine serum (FBS) and 1% penicillin/streptomycin (Thermo-Fischer) under standard conditions (37 °C, humidified atmosphere, 5% CO_2_). At 80% confluence, cells were trypsinized and replated on CaF_2_ coverslips (20 x 20 x 0.35 mm, Crystan, Poole, U.K.) in 35 mm sterile Petri dishes (Corning). Cells were allowed to adhere to the coverslips overnight prior to FA feeding.

### Palmitic acid labeling

Media was replaced with DMEM containing 4.5 g/L glucose and L-glutamine supplemented with 1% FBS, 1% penicillin/streptomycin and 2-4% of unlabeled or labeled PA conjugated to BSA (final concentrations of palmitic acid between 60-120 µM).

### Live cell imaging preparation

At the appropriate times after feeding, cells were rinsed twice with 3 mL of PBS (Corning) and mounted in PBS on a glass microscopy slide (VWR) with a 10 µm double-sided tape spacer (Nitto, San Diego, CA).

### Fluorescence staining

24 hours after PA feeding, live cells were stained using ER-ID^®^ Red assay kit (GRP CERTIFIED^®^) (Enzo Life Sciences, Farmingdale, NY) and LipiSpot^TM^ 488 Lipid Droplet Stain (Biotium, Fremont, CA) according to manufacturers’ guides. In brief, a staining solution containing 200 µL of 1X assay buffer provided by the ER-ID^®^ Red assay kit, 0.15 µL of ER-Red Detection Reagent, 0.15 µL of Hoechst 33342 Nuclear Stain, and 0.2 µL of LipiSpot^TM^ 488 Lipid Droplet Stain was prepared. Cells were rinsed twice with PBS, followed by dispensing the 200 µL of prepared staining solution on the coverslip. Samples were incubated for 30 minutes at 37° C protected from light, after which samples were gently rinsed with 200 µL of 1X assay buffer and mounted on a glass microscopy slide as described above.

### OPTIR data collection

All imaging was performed as previously described (*11*) on a mIRage-LS IR microscope (Photothermal Spectroscopy Corporation, Santa Barbara, CA) integrated with a four-module-pulsed quantum cascade laser (QCL) system (Daylight Solutions, San Diego, CA) with a tunable range from 932 cm^-1^ to 2348 cm^-1^. Brightfield optical images were collected using a low magnification 10X refractive objective with a working distance of 15 mm. Spectra and infrared images were collected in co-propagating mode using a 40X Schwarzchild objective with a working distance of 8 mm in transmission mode. Data was collected with an IR laser power of 20 % and a probe power in the range of 11 %. All spectra and images were collected using PTIR Studio 4.6. (Photothermal Spectroscopy Corporation). For hyperspectral images, a step size of 250 nm and one acquisition was used.

### Fluorescence data collection

Fluorescent images were collected on the mIRage-LS IR microscope using a Prime BSI Express sCMOS (Teledyne, Thousand Oaks, CA). Fluorescent cells were excited using a Sola Light Engine (Lumencor, Beaverton, OR) and a GFP filter cube (TLV-U-FF-GFP, Thorlabs, Newton, NJ), a BFP filter cube (TLV-U-FF-BFP, Thorlabs), or a MCHE filter cube (TLV-U-FF-MCHC, Thorlabs). Images were collected using the 50X objective at 10% LED intensity.

### Data analysis

Fluorescence and brightfield images were processed in Fiji (NIH, Bethesda, MD) (*67*) Spectra were analyzed in IGOR Pro 9 (Wavemetrics, Portland, OR). Live cell ratio images were generated in Python 3.10 in Colab (Google, Mountain View, CA) (*68*)

### Ratio image generation

Hyperspectral ratio images were corrected to remove contribution from the water band and protein amide I as well as the overlapping lipid C=O peaks using correction values. Correction values were calculated after performing a multipeak fit analysis in IGOR Pro 9 (**Fig. S10**). Fits were obtained for the lipid C=O peak at 1744 cm-1, the lipid C=O peak at 1734 cm-1 and the amide-I and water band at 1650 cm-1 over 15 cells in spectra where both lipid peaks were visibly present before multipeak fitting. **Table 1** shows the fit parameters.

**Table 1.** Fit parameters for overlapping lipid, water, and amide I peaks. Peak numbers, locations, types, and width for the amide I and water band, 1744 cm^-1^ lipid carbonyl peak, and 1734 cm^-1^ lipid carbonyl peak

| Peak | Location<br>[$\text{cm}^{-1}$] | Peak<br>Type | Width<br>[$\text{cm}^{-1}$] |
| --- | --- | --- | --- |
| 1 | 1643-1655 | Gaussian | 50-100 |
| 2 | 1734 | Voigt | unbounded |
| 3 | 1744 | Voigt | unbounded |

As lipid peaks 1734 cm^-1^ and 1744 cm^-1^ closely overlap, all correction factors and ratio images were generated by using intensities at 1750 cm^-1^ as a representative of the lipid band at 1744 cm^-1^, as there is significantly less contribution of 1734 cm^-1^ lipid band on the 1744 cm^-1^ band at 1750 cm^-1^. Correction factors for the amide I and water band were calculated by dividing the intensity of peak 1 at 1734 cm^-1^ or 1750 cm^-1^ by the intensity of peak 1 at 1655 cm^-1^. These values were calculated across all cells in which multipeak fitting was performed and averaged to create a water correction factors for lipid peaks 2 and 3, respectively.

Correction factors for the two lipid bands were needed to remove contribution of the lipid peaks from each other, as these closely overlap. The correction factor for the 1734 cm^-1^ lipid band was calculated by dividing the intensity of peak 2 at 1734 cm^-1^ by the intensity of peak 2 at 1750 cm^-1^ to get correction factor *c.* Similarly, the correction factor for the 1744 cm^-1^ lipid band was calculated by dividing the intensity of peak 1 at 17 cm^-1^ by the intensity of peak 1 at 1734 cm^-1^ to get correction factor *d*. Both correction factors were calculated across all cells in which multipeak fitting was performed and averaged to create correction factors for each lipid band.

Lipid bands 1734 cm^-1^ and 1744 cm^-1^ were corrected for the amide I/water band intensity using **Equation 1** and **Equation 2**, respectively:

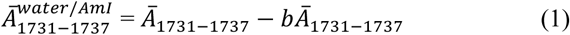

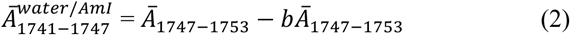

Where *Ā* is the average signal intensity at the indicated frequencies from full spectra of live cells and *Ā^water / Aml^* is the water and amide I corrected lipid band with the indicated frequencies. The variable *b* is the water and amide I correction factor, which is 0.22 for lipid band 1734 cm^-1^ and 0.15 for lipid band 1744 cm^-1^.

Lipid bands were then corrected for each other and the ratio image created using **Equation 3**:

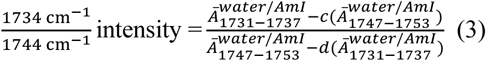

Where the variables *c* and *d* are the calculated correction factors discussed above and were 0.27 and 0.31, respectively. Live cell ratios were only performed in regions of the cell with significant lipid signal, defined as a 1744 cm^-1^ intensity of 0.01 or greater after vector normalization.

### Intermediate OPTIR spectra collection

The following intermediates were studied using OPTIR: tripalmitin, 1,2-dipalmitoyl-sn-glycerol, palmitoyl lysophosphatidic acid (Avanti Polar Lipids, Alabaster, AL), 1,2-dipalmitoyl-sn-glycero-3-phosphate (sodium salt) (Avanti Polar Lipids), and palmitic acid. Intermediates were dissolved at 25 mM in dichloromethane (Fisher Scientific) and chloroform.

### Lipid extraction and TLC

Cells were cultured in flasks and treated with 60 µM of either BSA, ^2^H oleic acid, or ^2^H palmitic acid. At 24 hours after treatment, cells were trypsinized and centrifuged at 3000 *g* for five minutes. The supernatant was discarded, and cells were resuspended in PBS. This process was repeated a second time, where cells were resuspended in 0.8 mL PBS. Lipids were isolated using the Bligh and Dyer method (*69*) In brief, a 3 mL of a cold chloroform/methanol (J.T. Baker, Phillipsburg, NJ) (1:2) solution was added to the resuspended cells, vortexed vigorously and incubated on ice for five minutes. 1 mL of chloroform and 1 mL of PBS was added and the solution vortexed again followed by centrifuging at 1000 *g* for 2 minutes. The bottom layer was recovered by a Pasteur pipette and a second extraction was performed on the cells by adding 1 mL of chloroform and centrifuging again at 1000 *g* for 2 minutes. The bottom layer was recovered again and dried under nitrogen, resuspended in 20 µL chloroform, and spotted on a silica gel plate. Lipids were separated using a mobile phase adapted from Chung *et al* (*70*) of hexane (Thermo Scientific): diethyl ether (Fisher Scientific): acetic acid (J.T. Baker) (80:20:1; v/v/v). TLC plates were then dried for 20 minutes and stained with a 0.2% solution of amido black (Thermo Scientific).

## Acknowledgments

The authors thank Jittima Weerachayaphorn at the Yale Liver Center for the Huh-7 cells and the Ellman group at Yale Chemistry for TLC materials.

## Author contributions

Each author’s contribution(s) to the paper should be listed (we suggest following the CRediT model with each CRediT role given its own line. No punctuation in the initials

Conceptualization: CMD

Methodology: HBC, CMD

Investigation: HBC, CMD

Visualization: HBC, CMD

Supervision: CMD

Writing—original draft: HBC, CMD

Writing—review & editing: HBC, CMD

## Data and materials availability

All data are available in the main text or the supplementary materials.

## Supplementary Materials

All data needed to evaluate the conclusions in this paper are present in the paper and/or the Supplementary Materials.

