## Supplemental Materials for "Optical photothermal infrared imaging of fatty acid esterification in the ER of living cells"

**This PDF file includes:**

Figs. S1 to S10

**Fig. S1. Full assignments of the OPTIR spectrum of 50 mM $^2$H PA in $CHCl_3$ (*71*, *72*).**

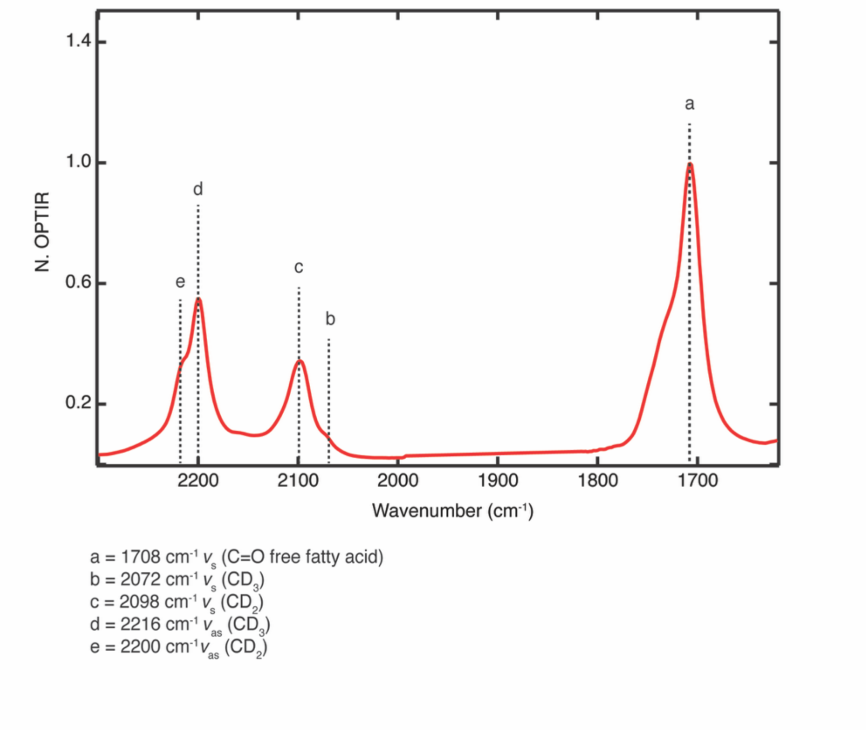

**Fig. S2. Detection limit of the OPTIR for the (A)** $v_{as}$ $CD_2$ stretch and **(B)** C=O carbonyl done by comparing spectra of 25 mM, 15 mM, 10 mM, 5 mM, 2 mM, and 1 mM of $^2H$ PA in ethanol. The detection limit for the $v_{as}$ $CD_2$ stretch was determined to be approximately 0.14 mM and the detection limit for the C=O carbonyl approximately 0.86 mM.

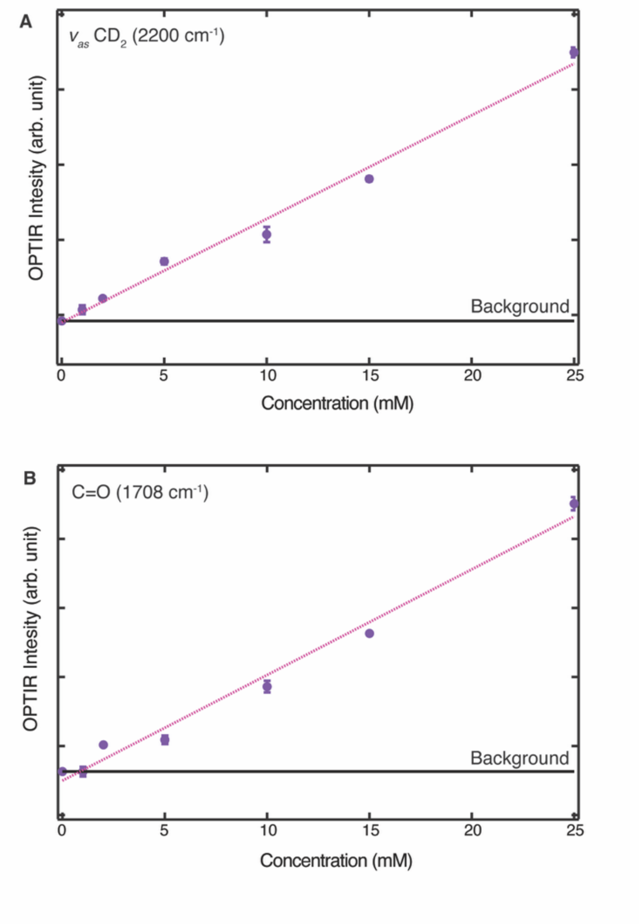

**Fig. S3. Representative spectra collected in a LD of a live Huh-7 cell 24 hours after feeding 60 µM of unlabeled PA conjugated to BSA at a 2:1 ratio.** The presence of a shoulder at 1734 $cm^{-1}$ with unlabeled PA is consistent with $^{2}H$ PA feeding.

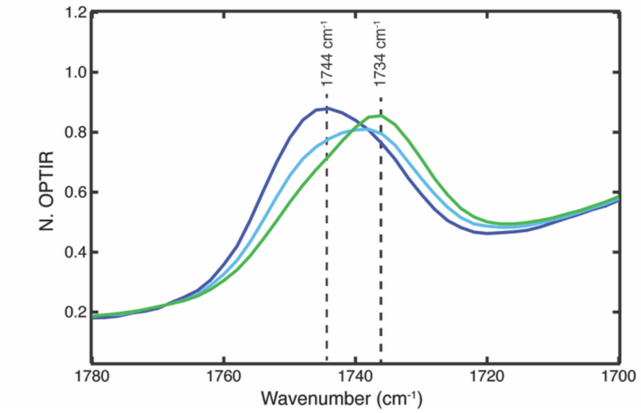

**Fig. S4. Irregularly shaped lipid droplets form when Huh-7 cells are fed PA but not OA or BSA.** Abnormal lipid droplets are apparent 39 hours **(A)** and 27 hours **(B)** after cells are fed 60 μM of $^2$H PA conjugated to BSA at a 2:1 ratio. Red arrows point out several oval shaped lipid droplets. Lipid droplets in cells fed 60 μM of $^2$H OA conjugated to BSA at a 2:1 ratio are spherical at the same time points, **(C)** 39 hours and **(D)** 27 hours. Lipid droplets in control cells fed BSA **(E)** and **(F)** are also regularly shaped. Scale bars are all 20 μm.

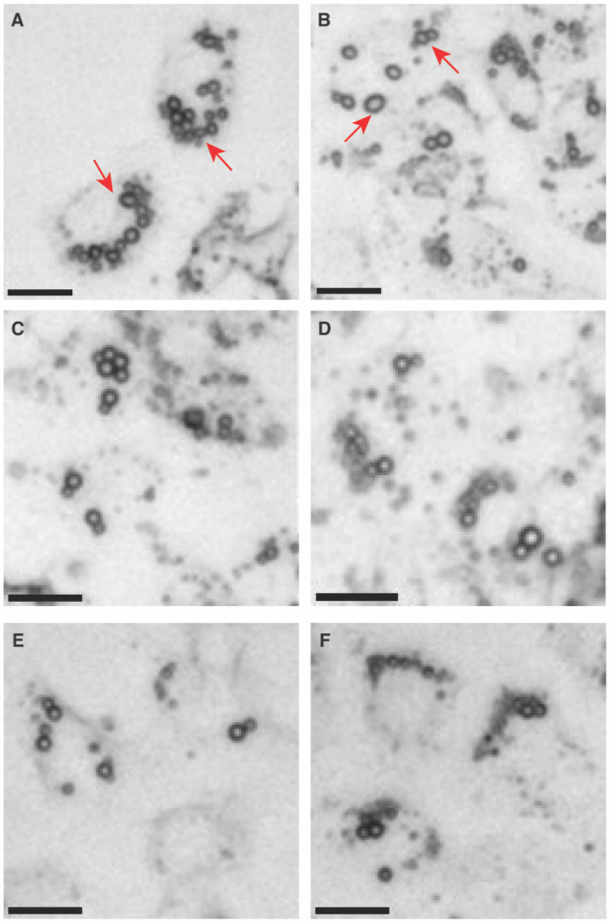

**Fig. S5. OPTIR spectra of 25 mM TAG precursors in DCM. Palmitic acid (red), 1,2 dipalmitin (blue), and tripalmitin (navy).**

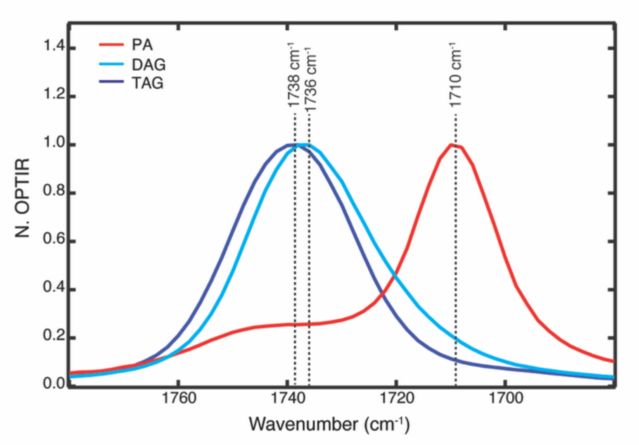

**Fig. S6. Cells fed 60 µM of $^2$H OA conjugated to BSA at a 2:1 ratio exhibit lipid carbonyl broadening and shoulders**. **(A)** Averaged normalized OPTIR spectrum collected across a LD in a Huh-7 cell 40 hours after feeding 60 µM of $^2$H OA. **(B)** A closeup of the lipid carbonyl region from 1700 cm$^{-1}$ to 1780 cm$^{-1}$.

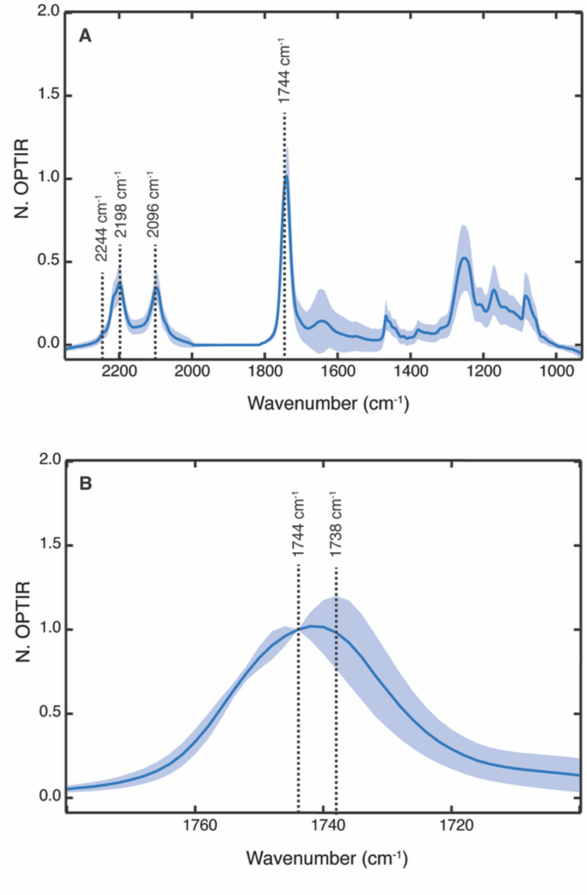

**Fig. S7. Average normalized OPTIR spectrum of a representative LD in a Huh-7 cell 24 hours after 120 µM of azido PA conjugated to BSA at a 2:1 ratio feeding.**

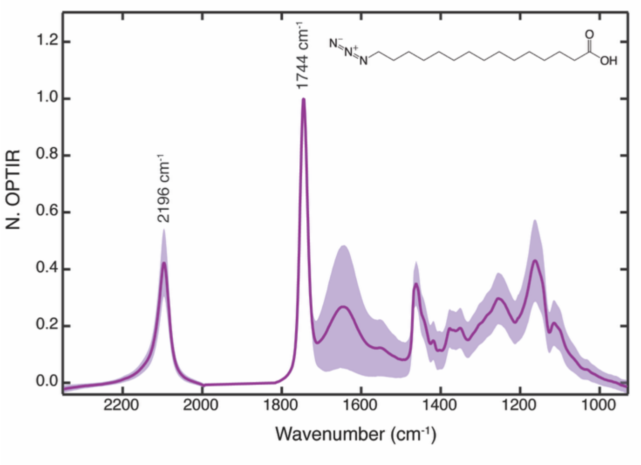

**Fig. S8. Comparison of the C-D region of Huh-7 cells fed 60 µM $^2$H OA conjugated to BSA at a 2:1 ratio.** **(A)** Spectra with (pink) and without (blue) strong 1738 cm$^{-1}$ intensity. A redshift of the asymmetric and symmetric $CD_2$ stretches is correlated with the appearance of a 1738 cm$^{-1}$ stretch alongside the 1744 cm$^{-1}$ lipid carbonyl. Enlarged view of the C-D region and its corresponding second derivative spectra is shown in **(B)** and **(C)**, respectively. Measurement of multiple cells and samples produced the same results. Spectra were normalized by prominent peak heights in the C-D region.

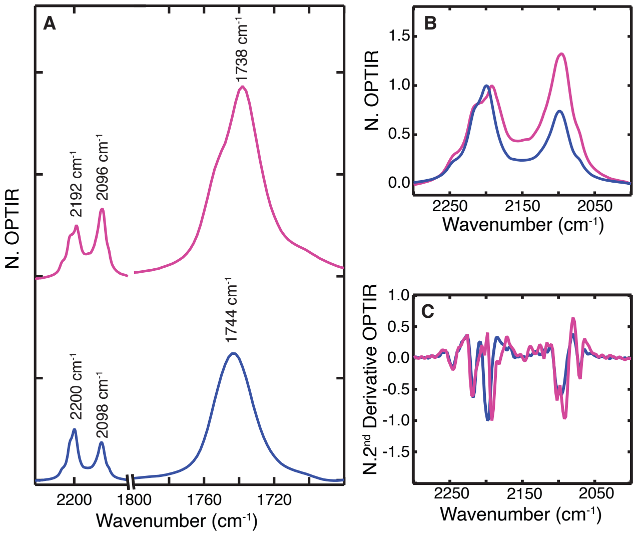

**Fig. S9. Rates of DNL decrease upon 60 µM $^{2}$H PA conjugated to BSA at a 2:1 ratio feeding (red) compared to control cells fed BSA only (purple).**

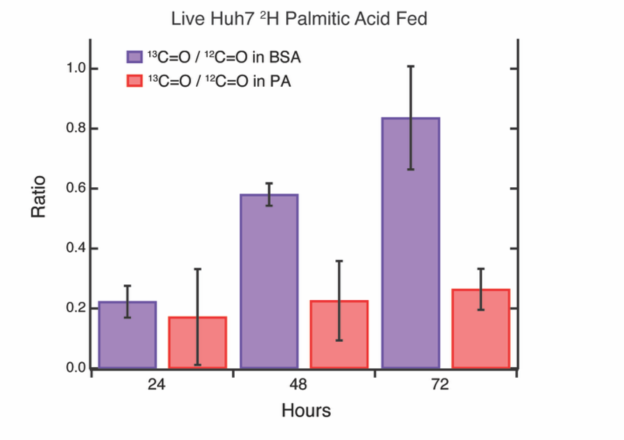

**Fig. S10. An example of the multipeak fitting performed in Igor where the two lipid peaks and the amide I/water band were fit to one Gaussian and two Voigt (cyan).** Residuals are shown on the top panel in pink.

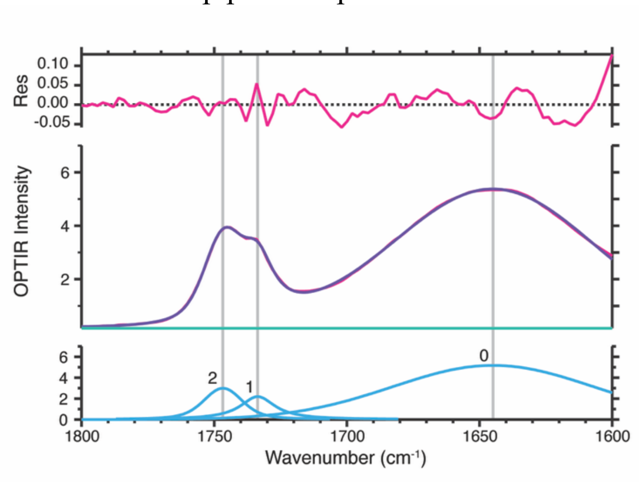
